# Performance comparison of high throughput single-cell RNA-Seq platforms in complex tissues

**DOI:** 10.1101/2023.04.04.535585

**Authors:** Yolanda Colino-Sanguino, Laura Rodriguez de la Fuente, Brian Gloss, Andrew M. K. Law, Kristina Handler, Marina Pajic, Robert Salomon, David Gallego-Ortega, Fatima Valdes-Mora

**Author notes:** Contributed equally (David Gallego-Ortega, Fatima Valdes-Mora).

## Abstract

Single-cell transcriptomics has emerged as the preferred tool to define cell identity through the analysis of gene expression signatures. However, there are limited studies that have comprehensively compared the performance of different scRNAseq systems in complex tissues. Here, we present a systematic comparison of two well-established high throughput 3’-scRNAseq platforms: 10x Chromium and BD Rhapsody, using tumours that present high cell diversity. Our experimental design includes both fresh and artificially damaged samples from the same tumours, which also provides a comparable dataset to examine their performance under challenging conditions. The performance metrics used in this study consist of gene sensitivity, mitochondrial content, reproducibility, clustering capabilities, cell type representation and ambient RNA contamination. These analyses showed that BD Rhapsody and 10x Chromium have similar gene sensitivity, while BD Rhapsody has the highest mitochondrial content. Interestingly, we found cell type detection biases between platforms, including a lower proportion of endothelial and myofibroblast cells in BD Rhapsody and lower gene sensitivity in granulocytes for 10x Chromium. Moreover, the source of the ambient noise was different between plate-based and droplet-based platforms. In conclusion, our reported platform differential performance should be considered for the selection of the scRNAseq method during the study experimental designs.

## Introduction

Single-cell RNA sequencing (scRNAseq) enables simultaneous profiling of gene expression of individual cells and is the tool of preference to define cell phenotypes and discover cell subsets and states ^1^. Single-cell transcriptomics provides an unprecedented resolution of the composition and functionality of cellular niches in homeostatic tissues and during disease ^2,3^. For example, scRNAseq has been used to unravel multicellular dynamic processes during embryogenesis and cell differentiation, tissue regeneration and morphogenesis, disease initiation and progression, and response to stimuli including drug treatments ^4–12^.

The ability to sequence genomic material combined with barcoding strategies to label each cell and RNA molecule extended the possibility of simultaneous analysis of barcoded cells in the same reaction, dramatically reducing cost and labour ^13,14^. The subsequent application of microfluidic devices to encapsulate individual cells in nano-droplet-sized bioreactors, like Drop-seq ^6,7^, or the development of high-density microwell plates to partition and capture individual cells, as Microwell-Seq ^15,16^ dramatically increased the scRNAseq scale, enabling the analysis of tens-of-thousands of cells in a single reaction. These high throughput scRNAseq methods can be technically challenging and difficult to control to deliver consistent outputs ^17^; however, commercial solutions have streamlined these processes, easing the technical demand and standardising procedures while ensuring consistent reagent quality, like 10x Chromium and BD Rhapsody ^18^.

At a first glance, all high throughput scRNAseq methods offer similar data outputs and they can interchangeably be used to interrogate any biological system. However, each method is technically different, with its own intrinsic capabilities, biases and limitations; thus, platform performance could vary with particular cell types and tissues, and specific platforms may be better suited to answer particular biological questions. There is limited information on a systematic comparison of the performance of different high throughput scRNAseq platforms and the available scRNAseq platform comparisons are based on relatively homogenous cell cultures ^19,20^, or artificial mixed pools ^21–23^. Thus, these comparisons are unable to assess each platform’s ability to distinguish cellular heterogeneity in a complex tissue scenario, failing to compare performance between cells of different lineages. For example, a recent study comparing five single cells/nucleus RNAseq high-throughput methods concluded that the commercial platform 10x Chromium was the top performing technology compared with the home brew methods ^24^. This study, however, did not included other commercial high-throughput platforms, such as the BD Rhapsody; and for the single-cell comparison, it only used cultured cells or PBMC samples which do not require tissue dissociation.

Here, we present a technical and biological comparison of two well-established and widely used 3’-scRNAseq commercial platforms, - 10x Chromium (10x Genomics) and BD Rhapsody (Becton Dickinson); using mammary gland tumours from the MMTV-PyMT mouse model, biologically complex but reproducible samples, ^25–28^. 10x Chromium is a droplet-based microfluidic platforms while BD Rhapsody uses microwell-based technology where cells are randomly deposited by gravity into an array of picoliter-size wells. All systems track the cell of origin with a cell barcode and count individual molecules using unique molecular identifiers (UMIs). Although these platforms are designed to produce similar readouts, essentially a digital count of genes in each cell, the device design (microfluidic or microwell), the nature of the capture beads, the molecular design of the barcodes and UMIs, and the essential nature of the molecular workflow for RNA copy and amplification strongly differs.

In this study, we evaluate how these platforms manage the challenges of complex tissues to produce meaningful data from scRNAseq and discover the strengths and weaknesses of each platform.

## Material and Methods

### Mouse model

The MMTV-Polyoma Middle T antigen (PyMT) was a gift from Dr. William J. Muller (McGill University) and its generation has been previously described ^26,27^. At ethical endpoint, (10% ± 3% tumour/body weight, which approximately corresponds to 14-week-old animals) the mice were euthanized, and size-matched tumours were harvested and processed for single cell digestions. Genotyping was performed at the Garvan Molecular Genetics facility (NATA accredited, ISO 17025) by PCR of DNA extracted from the mouse tail tip using the following primers: CGGCGGAGCGAGGAACTGAGGAGAG and TCAGAA GACTCGGCAGTCTTAGGCG. The touchdown PCR conditions were 94°C for 10 s of initial denaturation, followed by 10 cycles of 94°C 10 s, 65-55°C for 30 s and 72°C for 1 min and 10 s and then 31 cycles of 94°C 10 s, 55°C for 30 s and 72°C for 1 min and 10 s; the final extension is 72°C for 3 minutes. All animals used in this study are heterozygous for the PyMT gene.

All animal experiments were carried out according to guidelines contained within the NSW (Australia) Animal Research Act 1985, the NSW (Australia) Animal Research Regulation 2010 and the Australian code of practice for the care and use of animals for scientific purposes, (8th Edition 2013, National Health and Medical Research Council (Australia)). All experiments involving mice have been approved by the St. Vincent’s Campus Animal Research Committee AEC #17/03 and #19/02.

### Tissue digestion and single-cell isolation

PyMT tumours were digested as described in ^29,30^. Briefly, tumours were manually dissected into 3-5 mm pieces using surgical blades and further chopped to 100 µm using a tissue chopper. Samples were enzymatically digested with 15,000 U of collagenase (Sigma Aldrich Cat# C9891) and 5,000 U of hyaluronidase (Sigma Aldrich Cat# H3506) for 30 min at 37°C. The samples were then further digested by pipetting up and down with 0.25% trypsin (GIBCO Cat# 15090-046), in 1mM EGTA and 0.1 mg/ml of Polyvinyl alcohol dissolved in PBS for 1 min at 37°C in a waterbath. Red blood cells were then lysed with 0.8% ammonium chloride (Sigma Aldrich Cat# A9434) dissolved in water for 5 min at 37°C. Single cell suspensions were washed with PBS containing 2% FBS and spun at 200 x g for 5 min at 4°C between each step. The supernatant was aspirated and 1mg/mL DNase I (Roche Cat# 10104159001) was mixed with the sample before incubation with each step. Finally, cells were filtered through a 40 mM nylon mesh (Corning) and resuspended in PBS with 2% FBS. For the generation of low-quality-like samples, digested tumours were left overnight at 4°C, and check that viability was reduced by flow cytometry by at least 20% before processing the sample in auto macs.

All tumours were labelled with Annexin specific MACS beads using the Dead Cell Removal Kit (Miltenyi Biotec Cat# 130-090-101) following the manufacturers’ instructions and dead cells were removed by passing the labelled cells through the autoMACS® Pro (Miltenyi) (see more details at Salomon et al., 2019). High viable cells (> 85% viability assessed by DAPI in flow cytometry) were loaded into 10x Chromium, BD Rhapsody or the Drop-seq microfluidic device.

### Flow cytometry

The viability and cellular content of the main cell compartments of the tumours was assessed by flow cytometry. Digested tissue samples were washed with PBS supplemented with 2% FBS and centrifuged at 200 x g for 5 min at 4°C. The pellet was then resuspended with 2% FBS in PBS for use in flow cytometry analysis. For flow cytometry analysis, single cell suspensions were incubated with anti-human antibodies for EpCAM (Clone 30-F11, BD Bioscience, Cat# 561868) and CD45 (Clone HI30, BD Bioscience, Cat# 560566) on ice in the dark for 30 min before they were washed, centrifuged, and resuspended with 2% FBS in PBS for analysis using the BD FACSymphony^TM^ Cell Analyser. To check viability, cells were stained with DAPI in a 0.5μg/mL concentration (Invitrogen, D1306) at 4°C for 3 min immediately before running the samples in the flow cytometer. Flow cytometry data were analysed using the software package FlowJo (version 10.4.2).

### 10x Chromium

Cell viability was verified by microscopy with 0.4% of Trypan blue solution (Sigma-Aldrich, T8154) immediately before cells were loaded into the 10x Chromium Controller. We aimed to capture 8,000 cells with > 90% cell viability from each condition. Libraries were prepared using the Single Cell 3’ Library Kit V3 from 10x Genomics. Sequencing was performed on an Illumina NovaSeq 5000 using a NovaSeq S4 200 cycles kit (Illumina, as follows: 28bp for Read 1, 91bp for Read 2 and 8bp for Index) to an estimated depth of 20,000-30,000 reads per cell. Cell Ranger pipeline v3.0.1 was used for Fastq file generation, alignment to the mm10 transcriptome reference and UMI counting. Barcodes corresponding to empty droplets were excluded using cell calling algorithm from Cell Ranger based on EmptyDrops ^31^.

### BD Rhapsody

We adopted the MULTI-seq protocol ^32^ based on lipid-modified oligos (LMOs) for multiplexing different samples in the same BD Rhapsody cartridge. Single-cell suspensions from four different tumours were washed twice with PBS and incubated in a 200nM solution containing equal amounts of anchor LMO and sample barcode oligonucleotides (generously gifted by Prof. Gartner laboratory) on ice for 5min. After LMO-barcode labelling, we incubated the co-anchor LMO (final concentration of 200nM) in each sample for 5min on ice. 1mL of 1% BSA was added in each sample to quench the LMO binding and wash once with 1% BSA. The tumour samples were pooled in equal proportions, washed twice with 1% BSA and resuspended in the cold sample buffer (BD Rhapsody Cat. No. 650000062).

Cell viability was verified (>90%) by microscopy with 0.4% of Trypan blue solution (Sigma-Aldrich, T8154). Then, immediately before cells were loaded into a BD Rhapsody cartridge, viability was corroborated by Calcein AM (Thermo Fisher Scientific, Cat# C1430) and Draq7TM (Cat# 564904) in the BD Rhapsody™ Scanner. Cell capture, cDNA synthesis and exonuclease I treatment were performed using the BD Rhapsody Express Single-Cell Analysis System as per manufacturer’s instructions. Sequencing libraries of LMO tags and mRNA were prepared using a custom protocol combining BD Rhapsody’s mRNA whole transcriptome analysis (WTA) library preparation protocol and the MULTI-seq protocol ^32^. First, we performed 2 sequential random priming and extension (RPE) reactions of the capture beads with mRNA and LMO cDNA to increase assay sensitivity using half of the reagents for each round. Then, we used the RPE product from the supernatant to continue the WTA library preparation following BD’s workflow and the post-RPE beads were resuspended in a cold bead resuspension buffer to prepare for LMOs cDNA amplification. After three washes with the resuspension buffer, the beads were resuspended in 80μL of the MULTI-seq cDNA amplification mix containing 1μL of 2.5μM Multi-seq primer 5’- CTTGGCACCCGAGAATTCC-3’ and PCR was performed as follows: (95 °C, 3 min; 95 °C, 30 sec, 60 °C, 1 min and 72 °C, 1 min for 14 cycles; 72°C, 2 min; 4 °C hold). Next, we purified the PCR products of the LMOs using a 0.6x ratio of the SPRI beads and keeping the supernatant, followed by a left side selection using 1.8X ratio. Eluted DNA was then quantified using Qubit and the expected DNA size (∼95 to 115 bp) was confirmed with the Agilent 4200 TapeStation system before proceeding with the library preparation PCR (95 °C, 5 min; 98 °C, 15 min; 60 °C, 30 min; 72 °C, 30 min; 8 cycles; 72 °C, 1 min; 4 °C hold). Each PCR reaction contained 26.25 μl 2× KAPA HiFi HotStart master mix (Roche), 2.5 μl of 10 μM TruSeq RPIX primer (Illumina), 2.5 μl of 10 μM TruSeq Universal Adapter primer (Illumina), 3.5 ng barcode cDNA and nuclease-free water to a total volume of 50 μl. A final cleanup using 1.6x SPRI bead ratio was performed before sequencing. WTA libraries and LMO libraries were pooled and sequenced on an Illumina NovaSeq 6000 using a NovaSeq S1 2x100 cycles kit (Illumina, as follows: 100bp for Read 1, 100bp for Read 2 and 8bp for Index) to an estimated depth of 25,000 reads per cell for the WTA library and 2,500 reads per cell for the LMOs.

Fastq files from BD Rhapsody samples were processed using BD Rhapsody™ WTA Analysis Pipeline, v1.9.1. WTA reads were aligned to the mm10 reference genome and LMO sequences (8bp) were included as sample tags, allowing a maximum of 1 mismatch. The sample multiplexing option of the WTA Analysis pipeline was used to determine the sample of origin of each cell and singlets were used for downstream analysis.

### Single-cell data integration, clustering, and annotation

Single-cell clustering and annotation were performed using Seurat v5.0.1. A Seurat object of each sample was created using *CreateSeuratObject()* function without a minimum gene or counts filtering. First, samples from the same platform were merged and normalized and variable genes were detected using the SCTransform method ^33^. Cell clustering was determined by shared nearest neighbour (SNN) modularity using 15 dimensions and a resolution of 0.3 and dimensional reduction was performed using *RunUMAP()* using 15 dimensions in all three platforms. Secondly, data from both platforms were integrated utilizing either reciprocal PCA (RPCA), canonical correlation analysis (CCA), Harmony ^34^ and Join PCS to identify anchors. RPCA integrated object with 5 anchors and normalized using SCTransform were used for further analysis. Cell identities were annotated using SingleR signatures based on the mouse cell type reference generated by the Immunologic Genome Project ^35^ and based on single cell reference mapping using the *TransferData()* function and the previously characterized PyMT tumour dataset as reference ^30^. Bar plots for comparing cluster abundance across platforms were generated using *dittoSeq* package and dotplots and boxplots using *ggplot2*. Gene markers of specific cell types or clusters were identified using *FindMarkers()* or *FindAllMarkers()* functions from Seurat using logfc.threshold = 1, min.pct = 0.3 and logfc.threshold = 0.25, min.pct = 0.25, respectively. To determine gene sets enriched in the upregulated genes of a specific cluster we used the *enrichGO()* function from the *clusterProfiler* package, p-value was adjusted using Benjamini-Hochberg formula and the q value cutoff set at 0.2. Gene expression correlation between different samples was calculated using the *DESeq2* package. Cell communication analysis was performed using the *CellChat* R package ^36^.

### Public data processing and analysis

10x Chromium and BD Rhapsody from bone marrow and whole blood gene expression and antibody matrices (BioProject ID: PRJNA734283) were downloaded and re-processed using Seurat v4.2.1. Filtered 10x cells based on the default settings of Cell Ranger or based on a custom threshold of Protein UMI counts per cell (>10) as Qi et al. ^37^ were first demultiplexed using the Hashtag oligo (HTO) libraries and the *HTODemux()* function from Seurat. Barcodes with more than one hashtag were discarded. Cells from the bone marrow and whole blood in each platform were merged, clustered, and visualised as explained above.

### Ambient noise detection

The single-cell Ambient Remover (scAR) model was used to determine the gene expression profile of the empty droplets or wells representing the ambient noise, the ratio of ambient noise in each cell and to create a denoised matrix ^38^. For the mouse datasets, for the setup_anndata() function, the whole list of barcodes selected were used as the unfiltered matrix and the putative cells based on either Cell Ranger of the BD Rhapsody pipeline for the filtered matrix, using a probability threshold of 0.995. Empty barcodes were used to calculate the ambient profile and train the model. For the human dataset from BD Rhapsody the noise ratio was calculated the same way than the mouse dataset for both the Gene expression and protein matrices. For the human datasets generated in 10x Chromium, the putative cells used in the *setup_anndata()* function were based on the manually filtering the UMI Protein counts (>10).

### Statistical analysis

scRNAseq data was generated from 16 different tumours across 4 different mice. Specific statistical details for sample size and the threshold for statistical significance can be found either in the figure legend or in the method section. The data in bar graphs are presented as mean ± SEM, and the statistical analysis employed an unpaired t-test. Box plots display data by depicting the minimum and maximum values as whiskers, the median of the lower half (quartile 1), and the median of the upper half (quartile 3) as a box. The graphs and statistical analyses for scRNAseq were performed using R and for flow cytometry using GraphPad Prism.

## Results

### Experimental design and data processing

Our comparison is achieved by using PyMT tumours that originated by the oncogenic PyMT protein in a genetically controlled congenic mouse strain (FVBn). These tumours present multiple lineages and high cell diversity while retaining reproducibility due to the limited variability ^30,39,40^. In addition, tumours are biologically challenging samples, as they are not homeostatic tissues, and present tissue damage, hypoxia or cell death; features that will put to the test any scRNAseq method. Tumour chunks from the 4 different mice (4 tumours per mouse) were randomized and were digested and enriched for viable cells using Magnetic Associated Cell Separation (MACS) as previously described ^29^ (**Figure 1A**). Single cell suspensions containing viable cells (>90%) were loaded into a 10x Chromium, or BD Rhapsody chip (**Table 1** and **Figure S1A**) and run following the corresponding standard molecular workflow.

**Figure 1:**
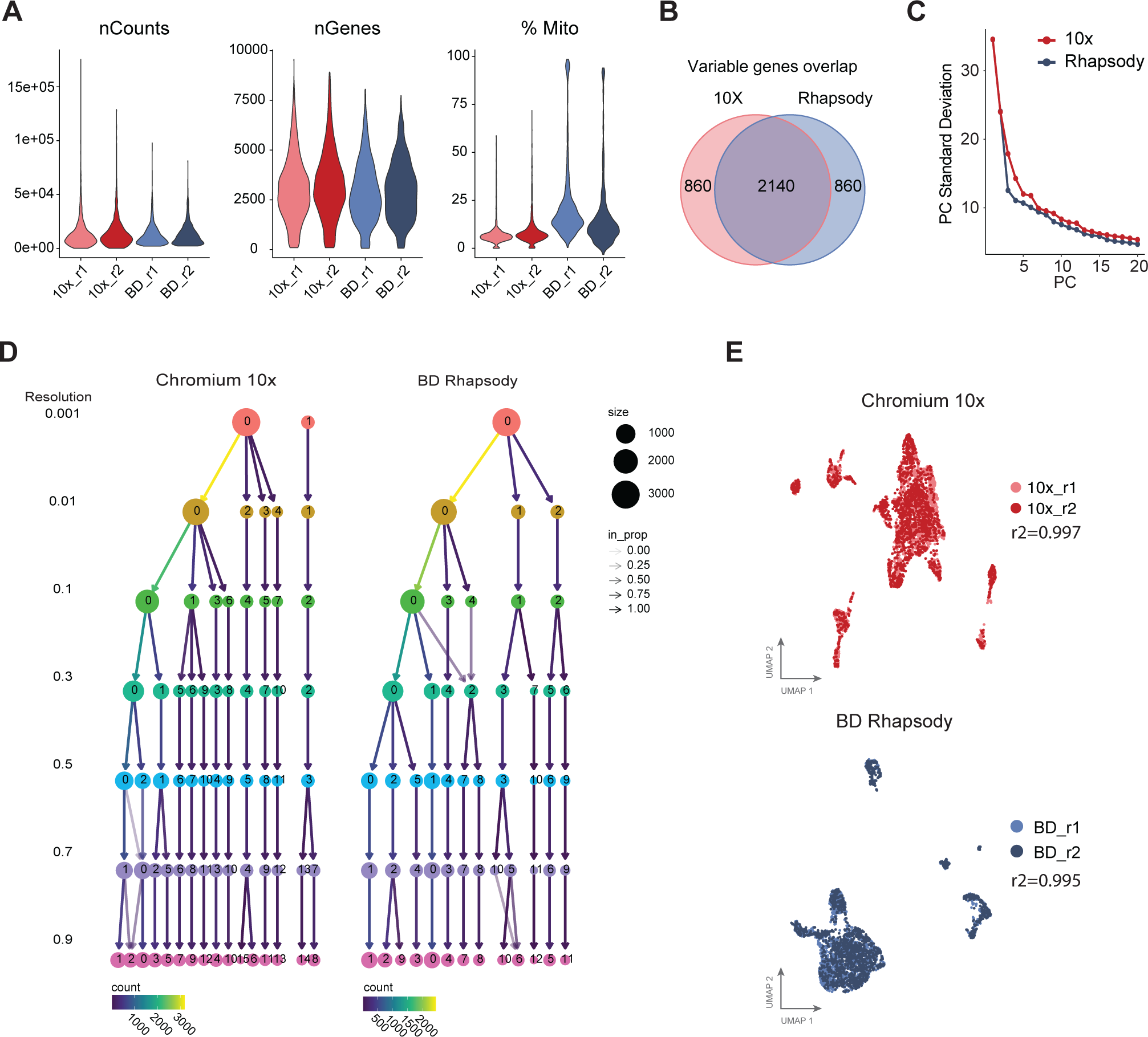
Measurements of the technical quality, sensitivity and reproducibility among scRNAseq platforms. A. Violin plots showing the number of UMIs (nCounts) and genes (nGenes) detected per cell, as well as the percentage of mitochondrial content (%Mito) per cell in 10x Chromium (blue) and BD Rhapsody (red) each using two biological replicates (r1, r2). **B.** Overlap between the top 3000 most variable genes in each platform. **C**. Principal component (PC) plot showing the total variability identified in each system measured as the standard deviation at increasing PCs. **D.** Cluster tree showing the phylogenic relationship of clusters in each platform at increasing resolutions. **E.** UMAP plots showing the two replicates in each platform.

**Table 1.**
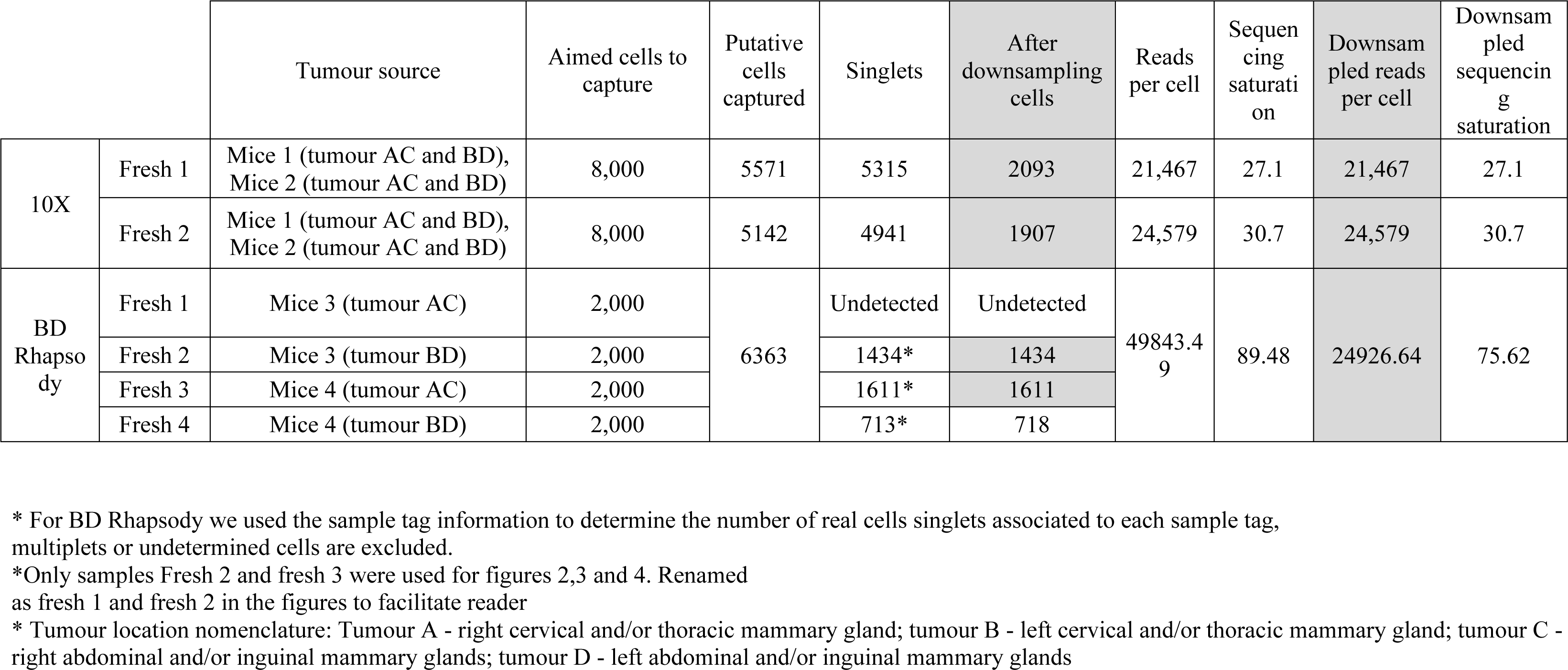

Each platform has a different read structure for barcodes and UMIs and each system has a different processing bioinformatic pipeline. Therefore, to replicate the standard user experience, we have used the intended pipeline for each platform to generate a Digital Gene Expression (DGE) matrix per cell: *cellranger* for 10x Chromium (10x Genomics) and the whole transcriptome analysis (WTA) pipeline in Seven Bridges Genomics cloud platform for BD Rhapsody. In all cases, data was subsequently integrated and visualised using the Seurat R package ^41^. A critical step for single-cell data processing is to determine how many real cell barcodes were detected. The latest version of the 10x Chromium pipeline, Cell Ranger 3.0, and BD’s pipeline in the Seven Bridges Genomics platform, now include a step that detects putative cells. Therefore, we filtered out empty barcodes based on the default parameters of each platform. Doublets were excluded using *DoubletFinder* ^42^ in the case of 10x Chromium, while for BD Rhapsody we used the information from the sample tags used to demultiplex tumour samples. After removing the empty droplets and doublets, we obtained 10,073 cells in 10x Chromium, and 6,363 for BD Rhapsody (**Table 1**). For BD Rhapsody we had 4 sample tags and after demultiplexing the two replicates with the greatest number of cells had 1,434 and 1,611 cells, therefore we subsampled the two replicates of Chromium 10x to obtain around 4,000 cells in total, which resulted in 2,093and 1,907 for downstream analysis (**Table 1**).

### Technical comparison: assessing sensitivity and reproducibility

An important quality metric for scRNA-seq is the sensitivity of gene and UMI detection per cell. This sensitivity metric was different between the platforms (**Figure 1A**); we detected a median of 2,995.5 and 2,791 genes, and 10,513.5 and 9,880 UMIs per cell in 10x Chromium and BD Rhapsody, respectively. Based on this analysis 10x Chromium and BD Rhapsody had a similar UMI and genes counts per cell at an equivalent number of reads per cell (**Table 1**). The proportion of mitochondrial genes is another metric commonly used to assess cell quality. Here we found that cells processed with 10x Chromium have significantly less mitochondrial content than BD Rhapsody samples, where 10x had a median of 6.5% of mitochondrial transcripts and BD Rhapsody 15.3% (**Figure 1A, right plot**).

An essential application for scRNAseq is to define single-cell phenotypes by their gene expression signature through dimensional reduction, principal component analysis (PCA), and clustering algorithms ^43^. We used the top 3,000 variable genes for dimensional reduction analysis in both platforms, noting that the variable genes detected in each platform were slightly different, with 2,140 (71.3%) common genes **(Figure 1B**). The total variability identified in the system, measured as the standard deviation at increasing principal components (PCs) had similar standard deviations per PC between platforms (**Figure 1C**). When we examined the clustering capabilities at increasing resolutions, we found that at the lowest resolution, only 10x Chromium was able to split the cells into two clusters suggesting that their detected transcriptome was more distinct, moreover, at increasing resolutions (>0.1), we found that 10x had a greater number of clusters and more consistent cluster composition than BD Rhapsody (**Figure 1D**). This consistency was measured by the capability of a cluster to further split into smaller clusters at higher resolutions without cells crossing into other cluster branches (**Figure 1D, crossing arrows**). In conclusion, BD Rhapsody and 10x Chromium had similar gene variability but 10x Chromium outperformed BD Rhapsody at consistently assigning cells to cluster at increasing resolutions.

Finally, to assess the reproducibility of each platform, we looked at the correlation of gene expression between replicates (**Figure 1E**). We found that both 10x Chromium and BD Rhapsody had over 0.99 correlation values confirming high reproducibility for both platforms.

### Cell type identification and lineage biases between platforms

To directly compare the performance of each platform to resolve tumour heterogeneity, we merged (**Figure S1B**) and integrated (**Figure 2A and S1C**) the data from the two platforms into one Seurat object. For the integration, we tested canonical Reciprocal PCA integration (RPCA), correlation analysis (CCA), Harmony Integration ^34^ and join PCS (**Figure S1C**). We found a similar level of integration using all four methods. RPCA integration has been reported to be less prone to overcorrection ^44^, therefore, we used this approach for the downstream analyses.

**Figure 2:**
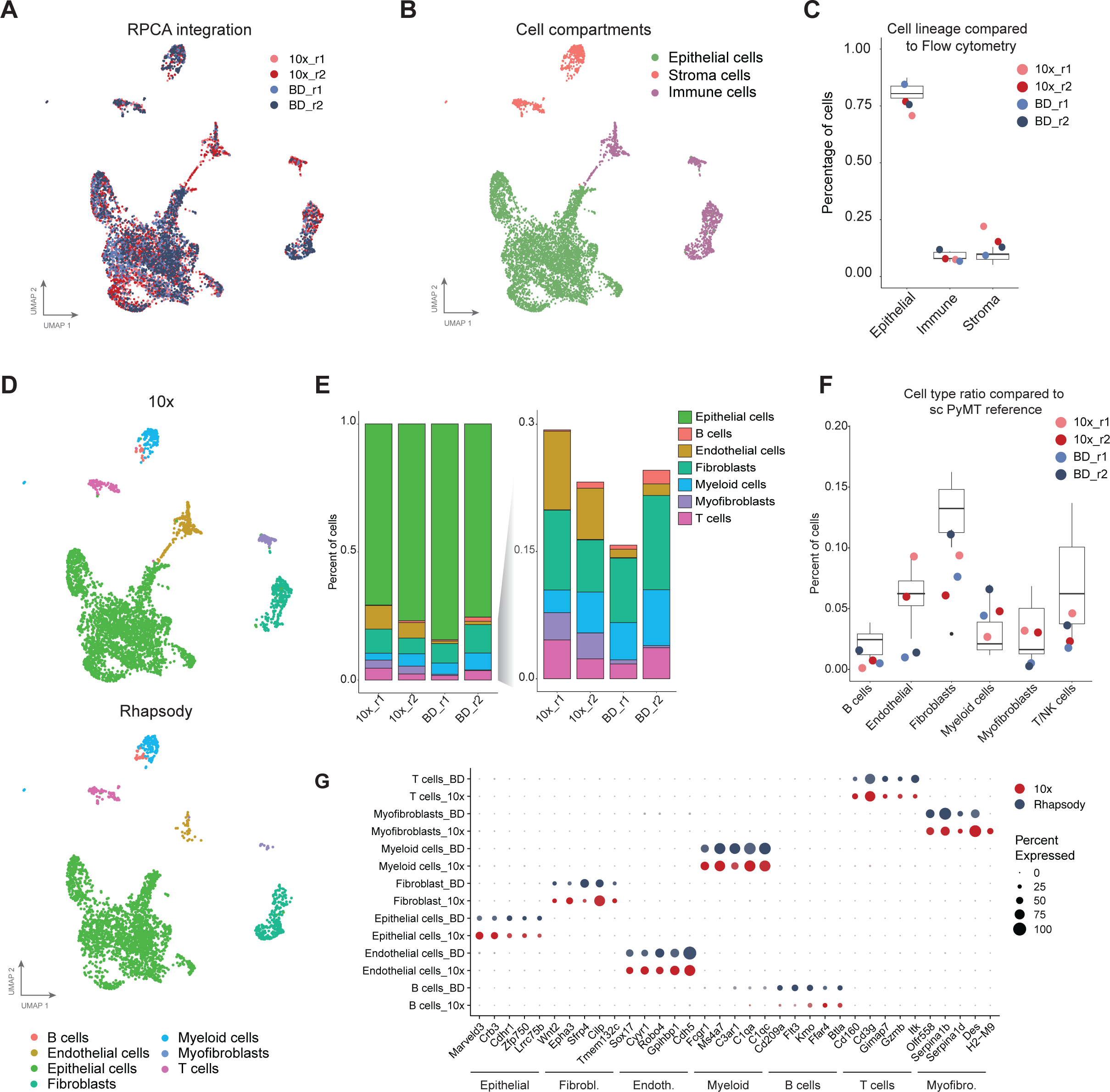
Identification of the main cell lineages in tumours by each scRNAseq platform. A. UMAP plot of all integrated samples using the RPCA method with 5 anchors. **B.** UMAP plot showing the three main cell lineages: epithelial, immune and stroma cells identified in all cells from all replicates and platforms after integration. **C**. Percentage of cells from each main cell lineage. The box plot represents the percentage of each cell lineage identified by flow cytometry and the colour dots represent each of our samples from the single cell dataset. **D.** UMAP plots showing the cell types assigned using label transfer analysis of the PyMT reference previously published ^30^, and split based on the platform. **E**. Bar plot of the percentage of cells from each cell type per sample (left) and excluding the epithelial cells (right). **F**. Cell type percentages of each cell type per sample. The box plot represents the percentages found in the PyMT reference ^30^ and the colour dots represent each of our samples from our single cell dataset. **G**. Dot plot showing the gene expression comparison of the top cell lineage marker genes between platforms.

For cell type identification, first, we divided the integrated object in the main cell lineages in PyMT tumours (epithelial, immune and stroma) **(Figure 2B)** and we compared their ratio in each platform to the observed percentage by flow cytometry, as it is the current gold standard method for cell type annotation and validation (**Figure 2C**). Overall, we found similar percentages between scRNAseq and flow cytometry, but the samples analysed through 10x Chromium had a slightly higher number, but consistent between replicates, of stroma cells and fewer epithelial cells. Next, we assigned each cell cluster to the main cell types based on SingleR ^35^ (**Figure S2A and B**) and also based on label transfer of the cell types from our previously published PyMT tumour reference ^30^ (**Figure S2C- S2E**). We found that all major cell types: B cells, T cells, myeloid cells, endothelial cells, epithelial cells, fibroblasts and myofibroblasts were detected in both platforms (**Figure 2D**), with similar percentages of cell types between replicates and platforms, except for endothelial cells and myofibroblasts (**Figure 2E**). BD Rhapsody identified a lower percentage of endothelial cells, from 1% of the total cells in BD Rhapsody compared to 9-6% in 10x Chromium, and myofibroblasts, 10 times less detected in BD Rhapsody compared to 10x Chromium. Consistently, the ratio of endothelial cells and myofibroblasts in BD Rhapsody deviated from the expected inter-tumour variability in the PyMT tumour reference atlas across eight tumours ^30^, suggesting that these differences were not a consequence of tumour heterogeneity (**Figure 2F)**. Of note, the percentage of fibroblasts was very variable between 10x/BD Rhapsody and the PyMT tumour reference atlas this could be due to a technical bias of the Drop-seq platform used in this atlas ^30^. We further confirmed the consistency of cell annotation across platforms by looking at the top markers of each cell type and confirmed that these markers were similarly expressed in all cells captured in both platforms (**Figure 2G**).

To determine how the different cell population ratios, gene marker dropout or ambient noise can affect downstream analysis, we performed cell-to-cell communication predictions using the CellChat software on these two platforms independently. We found that overall BD Rhapsody detected a larger number of interactions and higher interaction strength (**Figure 3A**). In line with the number of cells detected, BD Rhapsody detected fewer and weaker interactions between myofibroblasts with epithelial cells or fibroblasts and between endothelial and epithelial cells (**Figure 3B, red lines**), however, it detected more interactions between most of the cell line pairs and it had stronger interaction within the epithelial compartment, between the fibroblast and the epithelial cells and between myeloid cells and epithelial or fibroblast cells (**Figure 3B, blue lines).** CellChat analyses also revealed that some signalling genes were missing in BD Rhapsody such as PTN and OCLN in the epithelial cells, NT and SMA7 in fibroblasts and APELIN in endothelial cells, while 10x did not detect 16 genes related to interaction pathways including ITGAL, CD45 and BAFF from myeloid cells, CLEC, CD48 and CD137 from B cells or ICOS and CD86 from B cells (**Figure 3C**). Overall BD Rhapsody seems to detect more cell-cell interactions and as expected, changes in cell type proportions had an impact on the interaction strength detected with those cell types.

**Figure 3:**
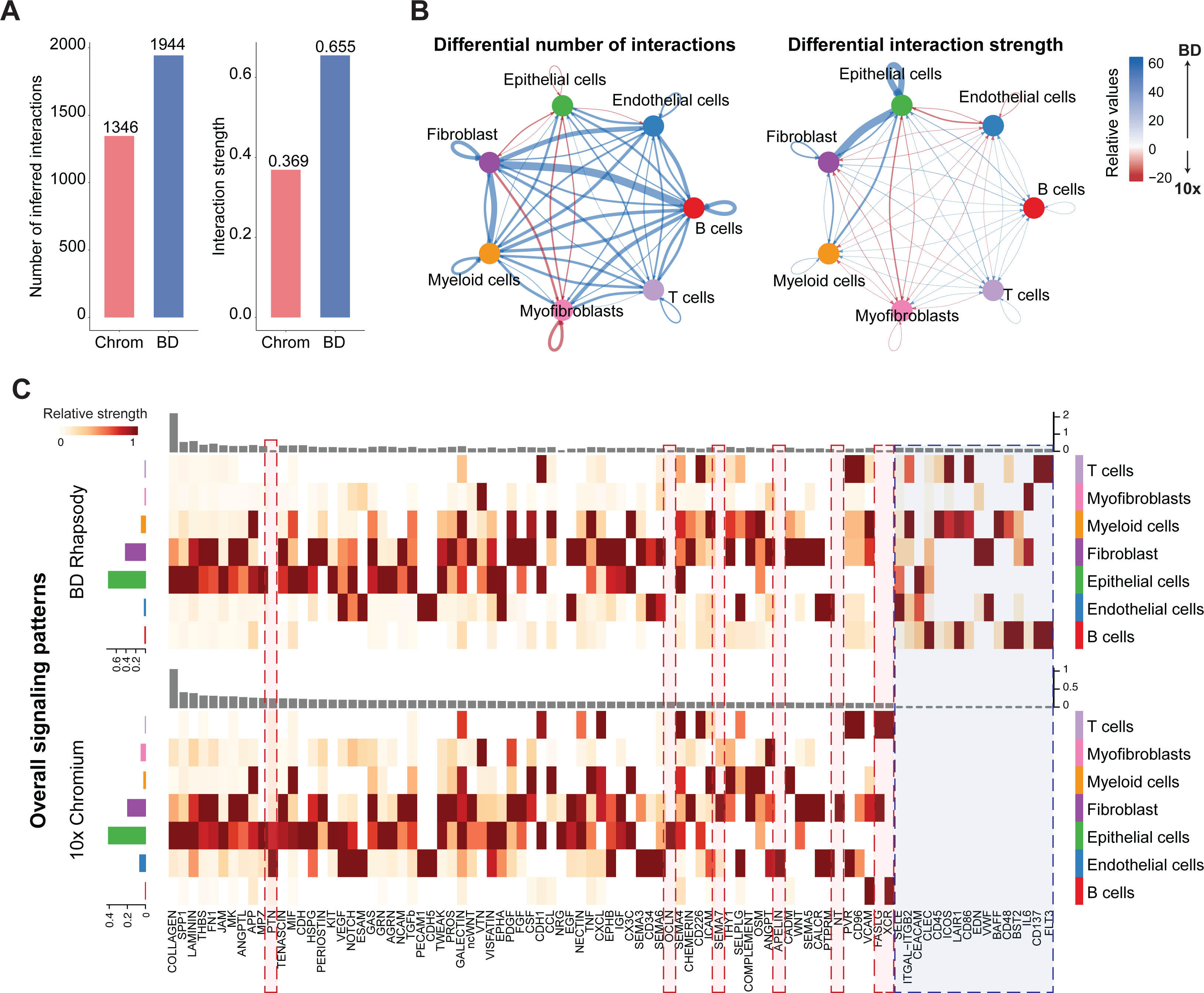
Comparison of cell-cell communication detection in each platform. A. Bar plot showing the overall number of interactions (left) and interaction strength in each platform (right). **B**. Circle plot visualizing the differential cell–cell interaction networks detected by each platform. In the left panel, the thickness of the line represents the number of differential interactions detected between the connecting cell types, while in the right panel, the width of the line corresponds to the differential strength of the interaction by assessing the level of expression of the ligand-receptor pairs after correcting the differences in cell type abundance. Blue lines represent higher number or stronger interactions in BD Rhapsody while red lines correspond to interactions enriched in 10x. **C**. Heatmap representing the relative signalling strength of the signalling pathways across platforms. The top grey bar plot shows the total signalling strength of each signalling pathway combining all cell types, and the coloured bar plot on the left shows the total signalling strength of each cell type combining all pathways.

In summary, 10x Chromium detected more endothelial cells and myofibroblasts, which resulted in different levels of cell-cell interactions detected with those cell types in each platform, however overall, the genes identify in BD Rhapsody per cell subtype were better on establishing cell to cell interactions, as a higher number and stronger interactions were detected.

### Cell subtype proportions across platforms

To further evaluate how each platform can distinguish among cellular subtypes within the main cell types in tumours, we performed subclustering of some of the main cell types. First, we selected the most abundant cell type in PyMT tumours, the epithelial compartment; unsupervised clustering revealed visual differential distribution in some cell clusters including clusters 1 and 3 out of the 7 clusters (**Figure 4A and S3A**). To infer the identity of each cell subcluster, we again used the PyMT tumour atlas ^30^ of the epithelial cell subtypes in the integrated epithelial cluster (**Figures S3B and S4C**), which resulted in the annotation of 8 epithelial subtypes manually (**Figure 4B**). As expected in this model, most cells were classified as luminal progenitors (LP), including a subset of hormone-sensitive luminal progenitor (LP-HS) cells where no major differences were observed between platforms (**Figure 4C**). Interestingly, the cell clusters where we visually observed differences (**Figure 4A**) were classified as basal (cluster 1) and luminal progenitor/ luminal hormone-sensing (LP/LP-HS) (cluster 3) (**Figure 4B and 4C**). Cluster 3, or LP/LP-HS cells, was missing from the PyMT reference (previously done using Drop-seq) and was heavily comprised of cells from the 10x Chromium platform. Gene set enrichment analysis identified that this Chromium-specific luminal progenitor cluster was enriched with genes involved in immune responses, including antiviral response and interferon (**Figure 4D)**. This suggests that the 10x Chromium platform detects more epithelial cells that are actively interacting with the immune system. Interestingly, basal cells (cluster 1) were virtually undetected in BD Rhapsody (3 cells detected), while in the 10x platform comprised around 2% of the epithelial compartment. We also identify differential percentage of cells between platforms in the Multi/Stem cluster (**Figure 4C**) with the highest proportion found in the BD Rhapsody replicates. Even though, the gene expression profile of the cluster 7 correlated with the multipotent/stem cell type (**Figure S3B** and **C**), we found that these clusters also had the highest percentage of mitochondrial gene content and the lowest number of genes detected in all platforms (**Figure 4E**), potentially suggesting damaged cells. Considering cells with high mitochondrial content but low number of genes detected as the definition of damaged cells, the 10x Chromium platform had the lowest ratio of damaged cells (**Figure 1A**), which may explain the absence of these cells and suggested that these are damaged cells.

**Figure 4:**
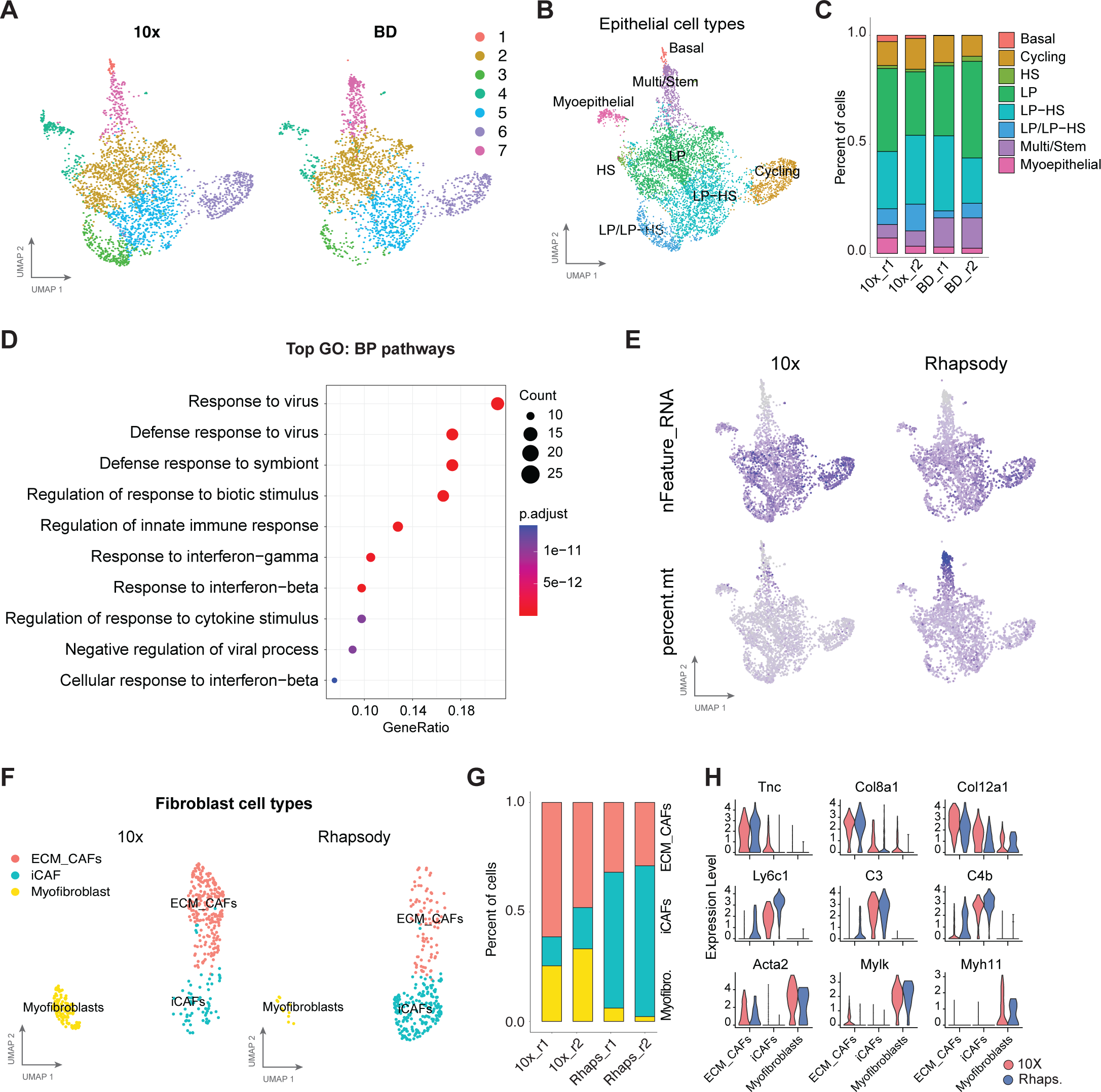
Cancer-epithelial and fibroblast subtype detection across platforms. A. UMAP projections of epithelial cells showing the unsupervised KNN clusters split by platform**. B** UMAP plot showing the epithelial cell types based on the annotated PyMT reference ^30^. (LP: luminal progenitor; HS: Hormone sensing) **C.** Bar plot of the percentage of cells from each epithelial cell type per sample. **D.** Gene set Enrichment analysis of the marker genes from cluster 3 (A) versus the rest of the luminal progenitor cells (GO: Gene Ontology. BP: Biological Processes). **E.** UMAP visualization of the number of genes (n_Feature_RNA) detected and the percentages of mitochondrial genes (percent.mt) per cell in each platform. **F.** UMAP projections showing the fibroblast subtypes divided per platform. (iCAF: immune cancer-associated fibroblasts; ECM_CAFs: extracellular remodelling cancer-associated fibroblasts). **G.** Bar plot representation of the percentage of each fibroblast subtype per sample**. H.** Violin plots for the expression of marker genes for ECM CAFs (top), iCAFs (middle) and myofibroblasts (bottom) comparing each fibroblast subtypes between 10x (red) and BD Rhapsody (blue) platforms.

Next, we did the same analysis in a less abundant population, fibroblasts. We again assigned cell types based on the fibroblast subsets in PyMT tumours ^30^ (**Figure 4F**) and found that within the fibroblast partitions, there were three clusters that correlated with either myofibroblasts or secretory cancer-associated fibroblasts (CAFs) which were subdivided into extracellular-matrix synthesis CAFs (ECM-CAFs) and inflammatory CAFs (iCAFs) (**Figure 4F**). The 10x Chromium platform detected a lower percentage of inflammatory CAFs, while the BD Rhapsody platform detected fewer ECM-CAFs and myofibroblasts (**Figure 4G**). In this context, the expression of key markers inflammatory CAFs, like Ly6c1 and C4b, was higher in BD Rhapsody, while myoepithelial markers, Acta2, Mylk and Myh11, were higher in the 10x data (**Figure 4H**).

Together these data suggest that, even though all major cell types were represented in all platforms, there are still cell type detection biases intrinsic to each platform which should be considered when analysing the tissue heterogeneity.

### Ambient noise comparison between 10x Chromium and BD Rhapsody in high and low-quality samples

One of the limitations of single-cell data is the technical noise caused by the ambient RNA contamination ^45^, amplification bias during library preparation and index swapping during sequencing ^46^. The nature of the cell partition method (nanowell or oil droplet) and the differences in the molecular workflows for RNA amplification in each platform (**Figure S1A**) could result in a differential origin of the technical and/or ambient noise. Ambient RNA is defined as the mRNA molecules that have been released from dead or stressed cells and are part of the cell suspension. The ambient noise is especially present in samples that require tissue digestion and in low-quality samples, such as tissues stored for an extended period before processing, or tissues with active cell dead or hypoxia regions like tumour tissues.

Our data suggest that noise might be handled differently by the commercial platforms (**Figure 1A, 4D** and **4E**). Thus, we studied how these two commercial platforms, performed with the technical noise from challenging samples. To recreate this “low quality sample”, we incubated PyMT digested tumours for 24h at 4°C. After 24h, cell viability dropped 20 per cent compared with the freshly digested sample as measured by DAPI positive cells in flow cytometry, indicating a substantial cell decay (**Figure 5A** and **Figure S4**). As done before with fresh PyMT tumour samples, we removed dead cells using MACS and processed the samples for scRNAseq using the 10x Chromium or BD Rhapsody platforms.

**Figure 5:**
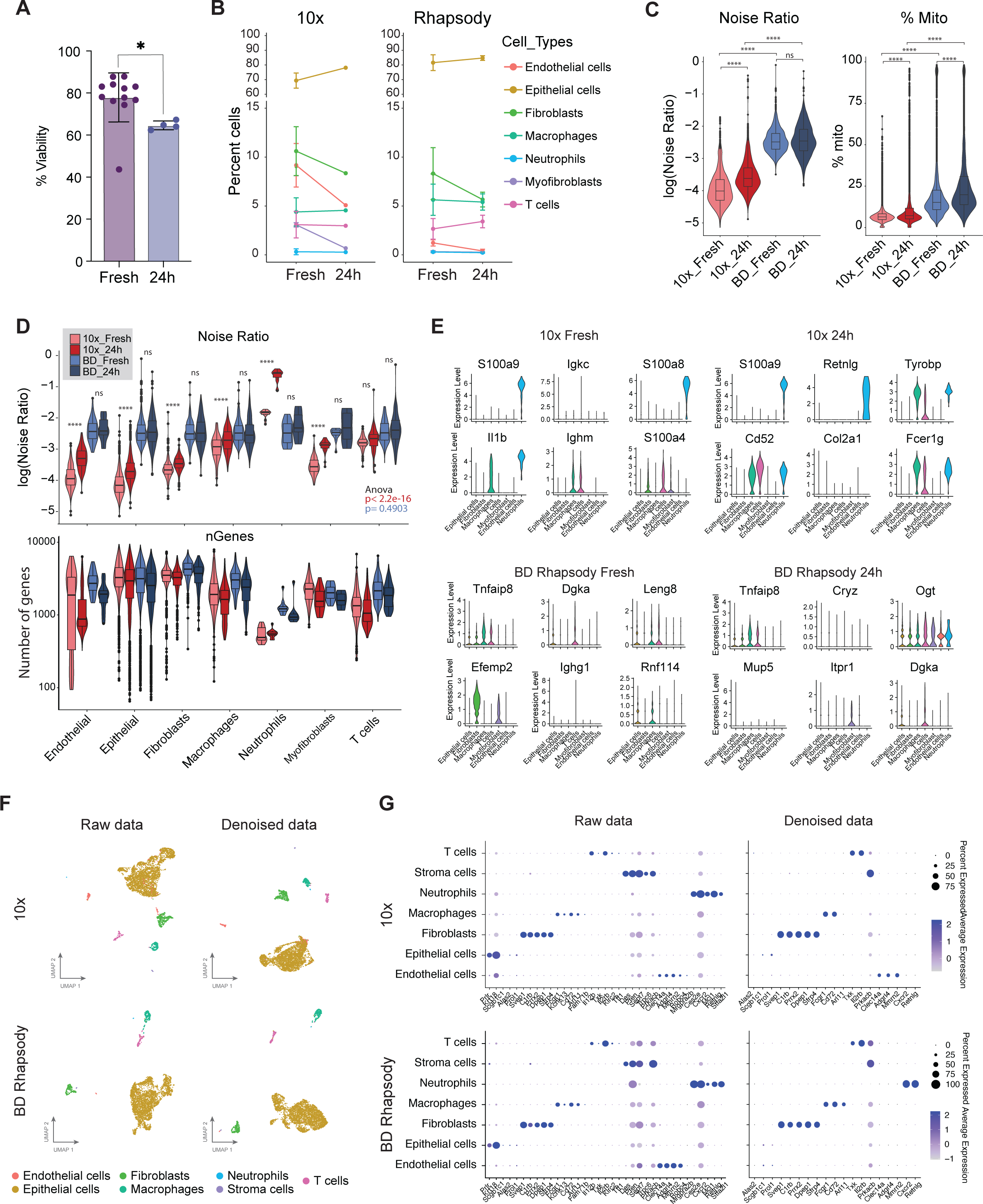
Analysis of the ambient noise in challenging samples in 10x Chromium and BD Rhapsody. A. Cell viability measured by flow cytometry as the percentage of DAPI negative cells in digested fresh tumours (Fresh) and simulated low-quality tumour samples (24h). **B.** Line plot showing the changes in percentage of cells for each cell type between fresh tumours and low-quality tumours (24h). **C.** Violin plots showing noise ratio (left) and percentage of mitochondrial content (% Mito, right) per cell in each condition. Statistical analysis of the comparisons between fresh and low-quality (24h) samples was performed using paired t-test adjusted with the Bonferroni method, ****adjusted p-value<0.001; ns = not significant. **D.** Violin plots split by cell type of the noise ratio (top) and the number of genes (bottom) detected in each cell. The y axis in the bottom panel is shown in logarithmic scale. Statistical analysis of the noise ratio comparisons per cell type between fresh and low quality (24h) samples in each platform was performed using paired t-test adjusted with the Bonferroni method, **** adjusted p-value <0.001; ns = not significant. One-way ANOVA was also performed to compare the noise ratio distribution across all cell types in the fresh samples in 10x Chromium (p-value=0.4903), or BD Rhapsody (p-value=2.2e-16). **E.** Violin plots for the gene expression levels of the top 6 most abundant genes in the ambient profile per cell type in each condition. **F**. UMAP projections using either the Fresh and 24h integrated raw data (left) or the denoised data (right) using scAR for 10x or BD Rhapsody. **G.** Dot plot showing average expression of the top gene markers of each cell type using the integrated raw data or denoised expression matrix.

First, we compared the tissue heterogeneity of the damaged samples to the fresh cells. As for this analysis, we increased the number of cells to ∼4000 cells per condition (**Table 2**), we were able to further split the myeloid cell compartment into neutrophils and macrophages. We found that endothelial cells, fibroblasts, stroma cells and neutrophils were consistently lost on the damaged sample regardless of the platform where myofibroblasts, fibroblasts and endothelial cells had the highest loss (**Figure 5B)**.

**Table 2.**
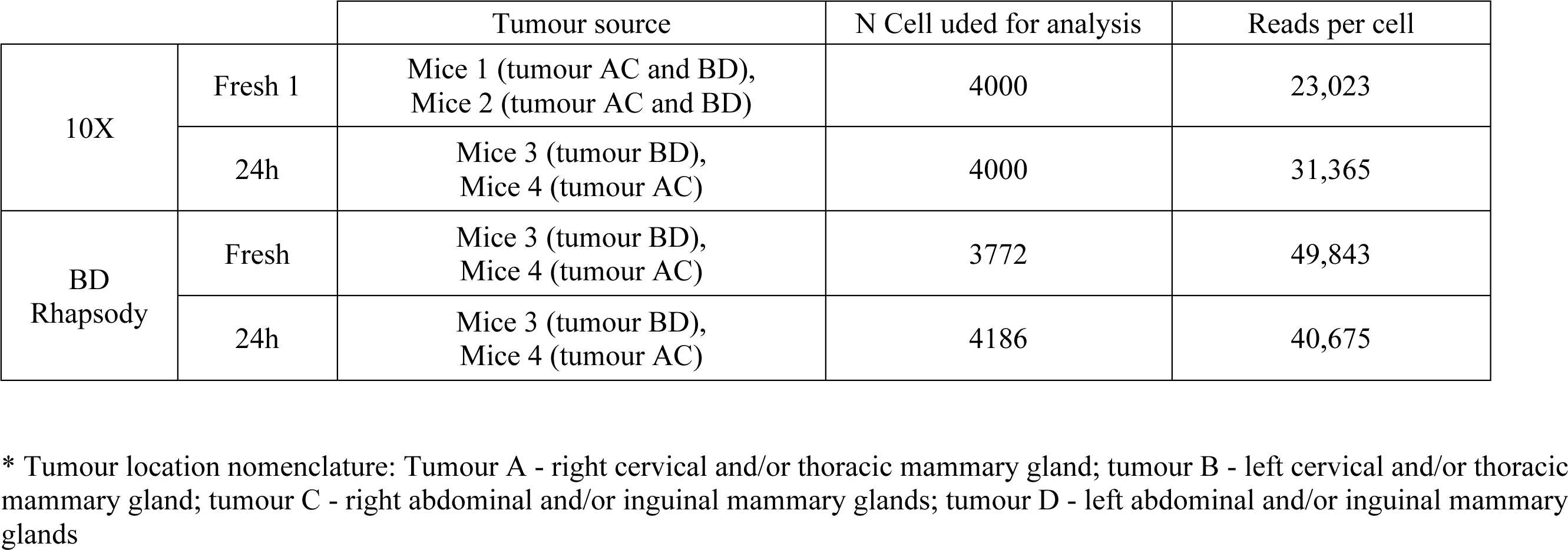

|  |  | Tumour source | N Cell used for analysis | Reads per cell |
| --- | --- | --- | --- | --- |
| 10X | Fresh 1 | Mice 1 (tumour AC and BD),<br>Mice 2 (tumour AC and BD) | 4000 | 23,023 |
|  | 24h | Mice 3 (tumour BD),<br>Mice 4 (tumour AC) | 4000 | 31,365 |
| BD<br>Rhapsody | Fresh | Mice 3 (tumour BD),<br>Mice 4 (tumour AC) | 3772 | 49,843 |
|  | 24h | Mice 3 (tumour BD),<br>Mice 4 (tumour AC) | 4186 | 40,675 |

\* Tumour location nomenclature: Tumour A - right cervical and/or thoracic mammary gland; tumour B - left cervical and/or thoracic mammary gland; tumour C - right abdominal and/or inguinal mammary glands; tumour D - left abdominal and/or inguinal mammary glands

Next, to assess the technical noise found in each platform, we used the single-cell Ambient Remover (scAR) method, a universal model to detect noise across single-cell platforms based on probabilistic deep learning of the ambient signal ^38^. Interestingly, around 5% of cells in both platforms and in any condition (fresh or damaged) were considered empty cells as their transcriptome was indistinguishable from the ambient signal of the empty barcodes (**Figure S5A**). Those so-called “empty cells” also had a high percentage of mitochondrial content (**Figure S5B**) and low gene content and therefore confirming that these assigned “cells” were instead damaged cells that thus needed to be removed from downstream analysis. When we compared the noise ratio of the remaining cells, we found that BD Rhapsody had significantly more ambient noise per cell than 10x Chromium (**Figure 5C**, 0h). However, the ambient contamination (noise ratio) was significantly increased in the damaged samples in the 10x Chromium but not in BD Rhapsody (**Figure 5C**, left plot). In addition, we confirmed our previous analyses of a higher percentage of mitochondrial content in Rhapsody (**Figure 1A**), and, as expected, in the damaged samples the mitochondrial content increased similarly in both platforms (**Figure 5C**, right plot). The number of genes found per cell decreased in the damaged samples compared to their fresh counterpart; however, the decrease on the number of genes was more pronounced with BD Rhapsody (**Figure S5C)**.

To evaluate which cell lineages are most affected by the technical noise in each platform, we compared the noise ratio per cell type and platform in each sample condition. We found that in BD Rhapsody all cell types had a similar level of noise, regardless of their damaged condition, while 10x Chromium’s noise was notably different between cell types and there were significant differences between fresh and damaged conditions (**Figure 5D** top). Particularly, in 10x Chromium, neutrophils had the highest percentage of noise which increased to an even higher ratio in the damaged samples. As expected, neutrophils also had the lowest number of genes detected especially in Chromium 10x (**Figure 5D** bottom), independently of sequencing depth (**Figure S6A**). To understand which cell types and genes are driving the ambient signal, we looked at expression across cell types of the top genes contributing to the ambient pool based on scAR analysis (**Figure 5E**). Interestingly, in the samples processed with 10x Chromium, 3 and 5 out of the top 6 most abundant genes in the ambient RNA pool in the fresh and the damaged sample, respectively, were neutrophils markers, despite the low number of neutrophils identified in the sample. In contrast, we did not find any strong expression specificity among the top genes in the ambient of the samples processed by BD Rhapsody, except for *Efemp2*, a fibroblast marker, in the fresh sample. Remarkably, in 10x Chromium, the ambient expression of the *S100a9* neutrophil marker was > 25-fold times higher than any other cell type marker and was even higher in the damaged samples, while in BD Rhapsody the ambient gene expression was more uniformly distributed and did not show major differences between fresh and damaged sample (**Figure S6B**). To confirm the neutrophil biases from 10x Chromium in the ambient profile, we correlated the average transcriptome of the single cells with the ambient gene expression and observed a higher presence of neutrophil marker genes in ambient RNA than in the single cells in 10x Chromium, but not in BD Rhapsody (**Figure S6C**).

The scAR algorithm can denoise the count matrix based on the ambient composition. When we used the denoised data to resolve cell type clusters, we found that clusters are more distinct using denoised data and there is less background expression of non-specific markers (**Figure 5F)**. As expected, the expression of the neutrophil markers was completely lost in the 10x Chromium data in this cell subtype (**Figure 5G**).

In conclusion, the differential molecular design, the microfluidic versus microwell format and the sample quality are sources of the ambient noise. In BD Rhapsody there is a generalised level of noise that is independent of the cell of origin and sample quality, while in 10x Chromium, ambient noise is cell type-specific and overrepresented by the neutrophil population.

### Cell type bias noise validation on public datasets

To determine if the cell type biases and technical noise differences between 10x Chromium and BD Rhapsody were universal, we assessed another two public data sets, human whole blood and bone marrow, where more neutrophils are expected, and both platforms were used ^37^. This data was processed using CITEseq (10x Chromium) and Ab-Seq + whole transcriptome analysis (BD Rhapsody) and therefore it also included 30 antibody-derived barcodes in both datasets. Additionally, hashtag oligos (HTO) were used in 10x Chromium platform to demultiplex samples.

We first compared the abundance of each cell type detected based on the antibody markers in both platforms using the default parameters for filtering established by Cell Ranger and the Rhapsody pipelines (**Figure 6A**). Interestingly, the whole blood and the bone marrow samples processed with 10x had virtually no granulocytes (neutrophils, basophils, or eosinophils) (**Figure 6B**). This is in line with previous reports of a low number of neutrophils in human datasets detected using this platform ^47,48^. Interestingly, if we filtered the 10x Chromium cell barcodes using the custom threshold based on the Protein UMI counts per cell as Qi et al. ^37^, we recovered 4,477 and 4,186 cells, in the bone marrow and whole blood samples, respectively, from which the majority were neutrophils (**Figure 6C**). In addition, using this custom filtering for all samples, resulted in very similar percentages of each cell type in the bone marrow and whole blood between the 10x and BD platforms (**Figure 6D**). However, even though the UMAP using the protein matrix was able to distinguish all major cell types in 10x Chromium (**Figure 6E**), the low gene sensitivity in the granulocyte population, with a median of fewer than 100 counts per cell (**Figure 6F**), did not manage to further resolve the granulocyte clusters at the transcriptome level (**Figure 6F, and Figure 6E** right panel). In fact, when we run the scAR algorithm using the threshold for filtering cells based on Protein UMI counts, we found that in 10x Chromium some granulocytes, especially eosinophils, still had a higher RNA noise ratio, although not as pronounced as in our mouse dataset in comparison with other cell types (**Figure 6G and 5D)**. Overall, the noise ratio from both the RNA and the protein data was more similar across cell types in BD Rhapsody than in 10x Chromium, confirming that in BD Rhapsody the ambient noise is not cell type specific (**Figure 6G**). In summary, 10x Chromium was unable to detect granulocytes in human samples based exclusively on their RNA content, and their low gene detection hinders the denoising algorithms to distinguish empty droplets from real cells with low UMI counts.

**Figure 6:**
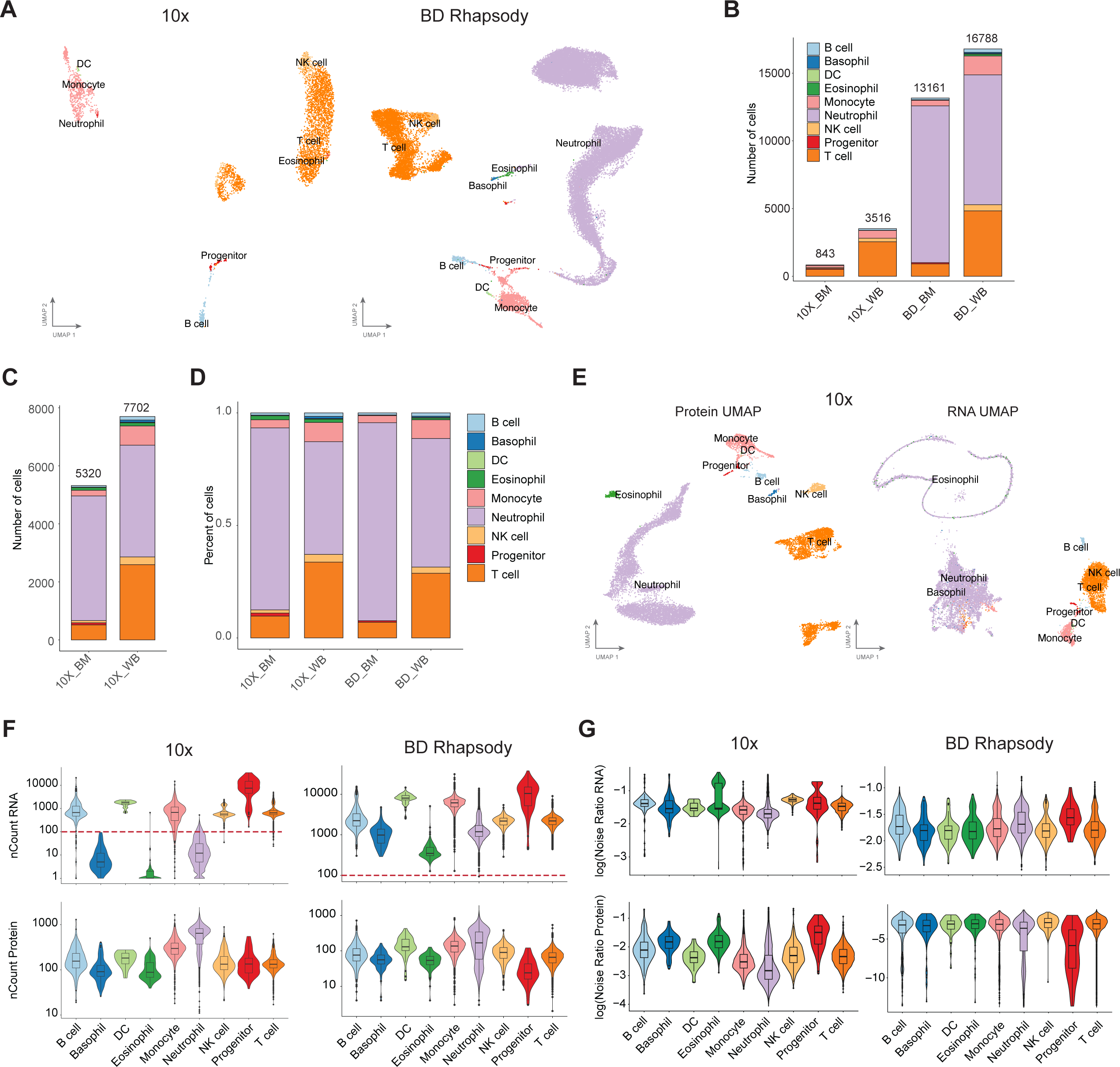
Performance evaluation for the detection of human granulocytes between 10x Chromium and BD Rhapsody. A. UMAPs of cell types found in human whole blood and bone marrow samples processed with 10x Chromium or BD Rhapsody using the transcriptome for dimension reduction. Cell barcodes were filtered using either Cell Ranger or BD Rhapsody pipelines. **B.** Bar plot showing the number of cells found in each cell type per condition. **C.** Bar plot showing the number of cells found in each cell type in 10x Chromium after custom filtering based on a minimum of 10 Protein UMI counts. **D.** Bar plot of the percentage of cells from each cell type per condition using the same custom threshold than panel C for 10x Chromium. **E.** UMAP plots of the human datasets processed with 10x Chromium and filtered based on custom threshold where the dimensional reduction was done using either the protein data (left) or the RNA data (right) **F.** Violin plots of the number of RNA and Protein UMI counts in each cell split by cell type. The red dotted line highlights a 100 counts threshold. The y axis is shown in logarithmic scale. **G.** Noise ratio in each cell using either from the RNA (top) or the protein (bottom) data matrix and split by cell type. (BM: bone marrow; WB: whole blood)

## Discussion

Single-cell RNA-seq allows the classification of different cell types based on the gene expression profile of individual cells. The rise of commercial instruments and kits has enabled the use of this methodology by the broad scientific community. However, there are still technical challenges during sample preparation, cell capture and library preparation ^49^, that can affect the bioinformatic readout and consequently, the biological interpretation driven from the data. Therefore, it is crucial to understand the advantages and limitations of each platform to best tailor the selection of a platform to the sample type and the biological question that wants to be answered, in addition to the costs and expertise level of the user.

Here, we have compared the most popular commercial platforms, the microfluidic-based system, 10x Chromium, and the microwell plate-based system, BD Rhapsody, using mammary gland tumours from the PyMT breast cancer mouse model ^25–28^. Analysing samples that require tissue digestion prior to the single-cell experiments is more challenging as the damaged cells can introduce additional ambient noise and different cell-type vulnerabilities can change the observed tissue heterogeneity. An advantage of our design is the comparison of tumour samples from a transgenic mouse model digested using the same protocol and in the same laboratory, which allows us to discard sample preparation as the source of variability. In addition, we have created low-quality samples using tumours from the same mouse model with reduced viability to assess how the quality of the sample affects the readout from 10x Chromium and BD Rhapsody. Frequently, the samples processed for scRNAseq have high levels of damaged cells, for example, tumour samples have necrotic regions or increased cell death due to their high cell turnover or exposure to treatments, such as radiotherapy or chemotherapy. Transportation of the biological material to the laboratory where cell capture is performed may also increase the cellular stress of the sample. In this study, we have used a biologically relevant tissue and replicated the standard user experience to perform a realistic assessment of the different technologies in challenging samples.

Our comprehensive data analysis identified, first, at the technical level, that 10x Chromium and BD Rhapsody had comparable overall gene sensitivity and reproducibility at similar sequencing depths. Next, we measured tissue heterogeneity and found a lower percentage of endothelial and myofibroblast cells in BD Rhapsody compared to 10x Chromium. However, comparing their ratios with the flow cytometry data, we found that 10x Chromium tends to detect more cells from the stroma than BD Rhapsody and flow cytometry. This suggests that 10x Chromium may enrich stroma cells which could be beneficial if you are interested in those populations. Interestingly, endothelial, and myofibroblast cells are reduced in low-quality samples in both platforms. We also found differences within the epithelial compartment, including an immune-responsive cluster that were enriched in the 10x Chromium samples. Orthogonal validation of this cell population is needed to understand its origin.

In the immune compartment, the biggest differences were found within the myeloid cells, especially in granulocytes, which 10x Chromium was not able to detect granulocytes neither in the PyMT mammary tumours nor in the human whole blood or bone marrow datasets, due to the low gene detection, which misclassifies them as empty droplets. It is known that granules in neutrophils, eosinophils and basophils have high levels of nucleases including RNases ^50^. This has been suggested to be the reason why granulocytes have lower genes and counts detected than other cell types ^37,51^. We have also confirmed in our PyMT dataset and human blood and bone marrow public datasets that granulocytes, including neutrophils, have the lowest number of genes detected. Interestingly, BD Rhapsody’s neutrophils had an average of 1,240 or 602 genes while in 10x Chromium neutrophils only had 551 or 10 in the mouse and human datasets, respectively. We hypothesise that there are two factors that may contribute to this phenomenon; firstly, neutrophils may be more sensitive to the pressure from the microfluidic devices which could cause their breakage and loss of RNA to the ambient pool in the 10x Chromium system; secondly, we hypothesise that the 10x platform is more sensitive to the presence of RNases released by neutrophils than BD Rhapsody. Supporting this theory, Qi et al found that cell doublets of neutrophils and T cells from 10x Chromium had less number of UMIs detected than T cell singlets, suggesting that the RNases from the neutrophils degraded the RNA of the T cells ^37^. The exact recipes of the lysis buffers used in 10x Chromium and BD Rhapsody are not disclosed, but it is possible that BD Rhapsody has stronger detergents or RNase inhibitors that degrade RNases. BD Rhapsody workflow also includes washes of the beads before cDNA generation which could remove any remaining RNases or strong detergent from the lysis buffer to not interfere with the reverse transcription reaction. Moreover, a recent report describing TAS-seq, a BD Rhapsody-based technique with improved cDNA amplification step, resulted in better cellular composition fidelity, especially in neutrophils, using their system compared to 10x Chromium and Smart-seq ^52^. Together, this demonstrates that BD Rhapsody is better suited for granulocytes research, and new strategies for cDNA amplification that increase the number of genes detected per cell will further improve the transcriptional resolution of this challenging cell population.

Mitochondrial content has been commonly used as a sign of broken cells, where their cytoplasmatic RNA has been lost but the RNA contained in the mitochondria remains ^53^. In fact, we see an increase in the mitochondrial fraction in the damaged samples. Surprisingly, all cells processed with the BD Rhapsody have a higher percentage of mitochondrial genes, even though they are not considered damaged. This phenomenon has also been recently reported by Salcher et al. ^54^. A possible explanation for this is that the lysis buffer of BD Rhapsody is more effective at digesting the organelles including mitochondria and therefore releasing the mitochondrial RNA into the cytosol to bind to the beads. For this reason, different thresholds of mitochondrial content should be used to filter out damaged cells in BD Rhapsody or 10x Chromium.

Ambient RNA release during the sample preparation and cell capture introduces noise to the data affecting the downstream analysis. Here, we have compared the level of ambient noise present in each barcode corresponding to putative cells and the gene expression of the ambient RNA from empty barcodes to identify vulnerabilities across platforms. We found that BD Rhapsody has a higher ratio of ambient RNA per cell than 10x Chromium, but the source of that RNA was coming from all cell types, suggesting a uniform contribution to the ambient pool. Ambient noise in BD Rhapsody could be created during the bead retrieval step where all beads are pooled and unbound RNAs could bind to free poly T tails from beads of different wells, and thus the likelihood of this occurring would be very similar across all cell types, in contrast to 10x Chromium where the RT occurs inside the droplet. Alternatively, the perceived high level of noise in BD Rhapsody could be due to the housekeeping genes’ contribution to the ambient RNA which are constitutively expressed in all cells at high levels. In contrast, the 10x Chromium ambient pool was biased towards certain cell types, in particular granulocyte, likely caused by damaged cells or RNases reducing the number of UMIs making it hard to distinguish between real or empty barcodes, as previously discussed. This suggests that BD Rhapsody may benefit from data-cleaning steps where the ambient fractions are removed, while 10x Chromium decontamination of the data may result in cell-type dropouts.

A limitation of our comparison analysis is the fact that we have used mouse mammary gland tumours, human whole blood and bone marrow only. Other tissues where different cell types are found may show other cell type biases, especially for lowly represented cell types. However, this study highlights the importance of using different technologies when assessing the population ratios of a sample and avoiding comparing cell type ratios across biological conditions performed in different platforms. Based on our analysis, we also found that the expected quality of the sample and origin of the tissue should also be considered when choosing a platform to perform single-cell RNAseq, as damaged tissues or tissues with high levels of RNases, such as bone marrow or spleen ^51^, would be more susceptible to RNA loss in 10x Chromium.

Complementary strategies, such as performing multi-omic studies where two or more molecular layers of information are investigated per cell ^55^, or spatial transcriptomics ^56^, as well as alternative methods for tissue preservation such as the ALTEN system ^11^ or parafilm fixation in combination with fixed RNA profiling ^51,57^, may also help dissect the true heterogeneity of complex tissues and overcome the current limitations of single-cell RNAseq.

## Data Availability

Data generated in this paper is available through Gene Expression Omnibus (GEO): GSE229765. PyMT mouse data processed with Dropseq is available in GSE158677. Human data sets were downloaded from PRJNA73428.

## Funding

This work is supported by Cancer Institute NSW Fellowship (2019/CDF002-CDF181218) to FVM and Cancer Institute NSW Fellowship (DG00625) and Cancer Council NSW project grant (RG18-03), NHMRC (2012941), NBCF Elaine Henry Fellowship (IIRS-21-096) to DGO; YCS is supported by Cure Brain Cancer Foundation (20210917)

## Acknowledgements

We thank the Garvan-Weizmann Centre for Cellular Genomics, the Molecular Genetics Facility, the Australian BioResources Pty Ltd facility and the KCCG Sequencing Laboratory from the Garvan Institute. We would also like to thank Dr Antoine de Weck for his guidance on the use of scAR and Prof Zev J. Gartner and Dr. Christopher S. McGinnis for gifting us the MULTI-seq kit reagents.

## Author contributions

Conceptualisation: DGO, FVM and YCS; Methodology: LRF, YCS, AMKL, KH; Formal analysis: YCS, BG; Investigation and validation: YCS, LRF, AMKL, KH; Resources: RS, MP, DGO, FVM; Data curation: YCS, DGO, FVM; Visualisation: YCS; Funding acquisition: FVM, DGO, MP, RS. Writing-Original, Reviewing & Editing: YCS, FVM, DGO. Supervision: FVM and DGO.

## Notes

### Competing Interest Statement

The authors have declared no competing interest.

### Summary of Updates

This manuscript version has been revised to reflect peer-reviewed feedback from reviewers. In particular, we have removed the Drop-seq data and comparison from the paper. We have also performed the comparative analyses between 10X and BD Rhapsody at a similar sequencing depth

