## Supplementary Figures for "Performance comparison of high throughput single-cell RNA-Seq platforms in complex tissues"

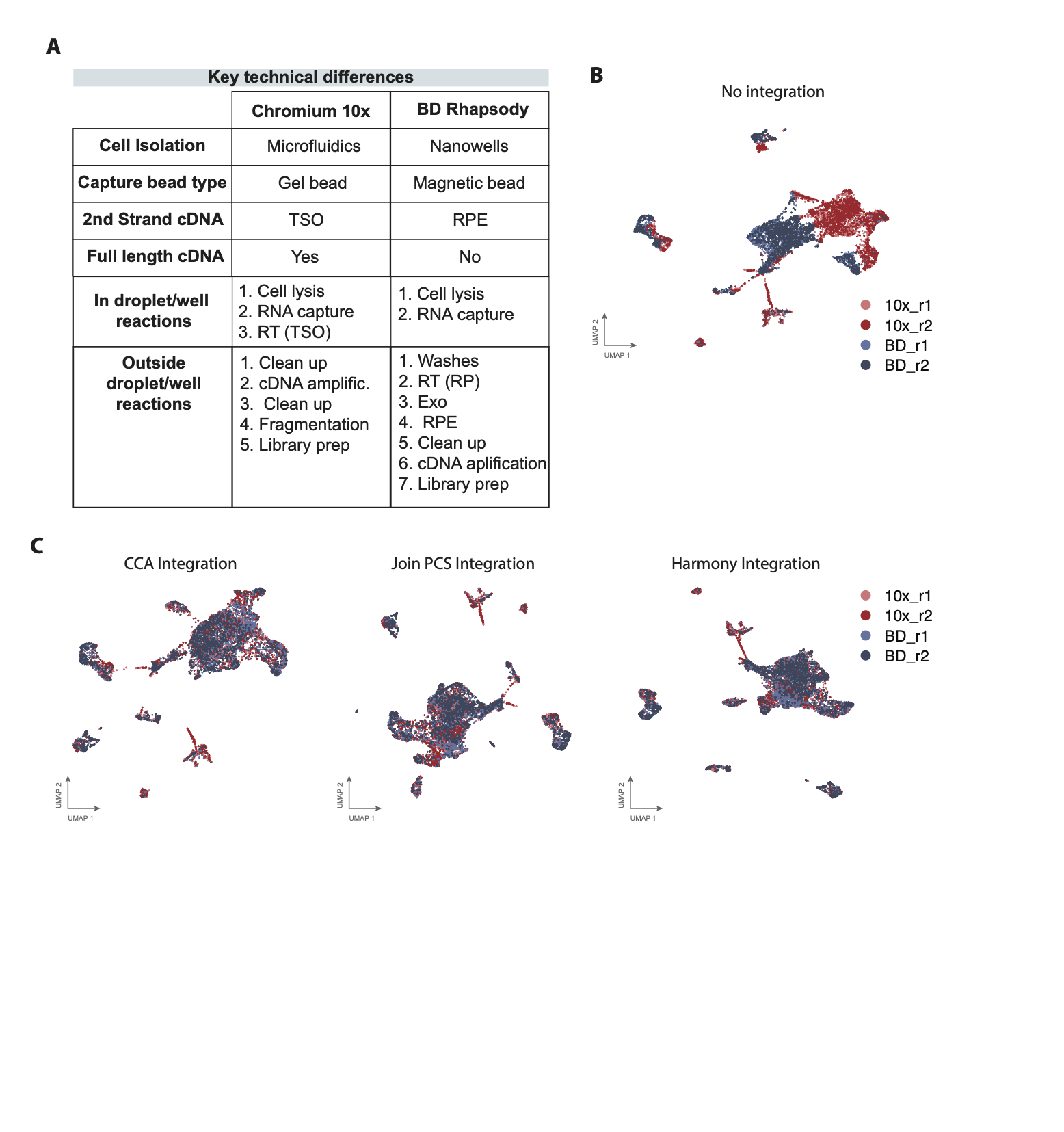

**Figure S1:** **A**. Table comparing the main difference among the three scRNAseq platforms used in this study. RPE: random priming and extension **B.** UMAP plot of non-integrated samples. **C.** UMAPs showing the integrated data across platforms using the Canonical Correction Analysis (CCA), the Join PCS Integration and the Harmony Integration methods.

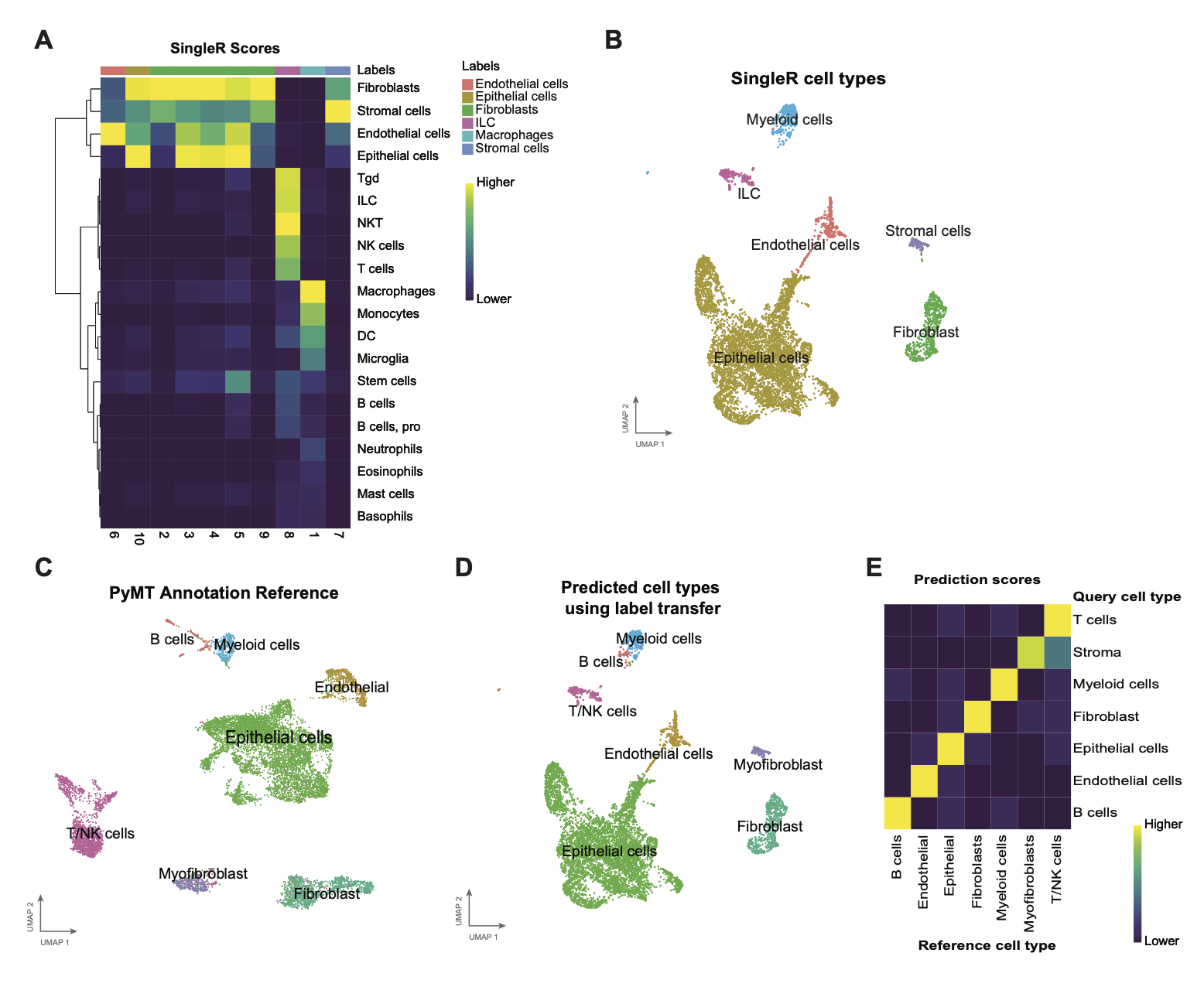

**Figure S2: A.** Heatmap showing the SingleR score for each cluster using ImmGenData Reference. **B.** UMAP plot of the assigned cell type labels based on SingleR analysis**. C.** Wild type PyMT annotation reference from GSE158677. **D.** UMAP overlaying the predicted cell types based on the label transfer method using the PyMT reference from panel C. **E**. Heatmap showing predicted scores for cell type from the PyMT reference on each single R annotated cell type.

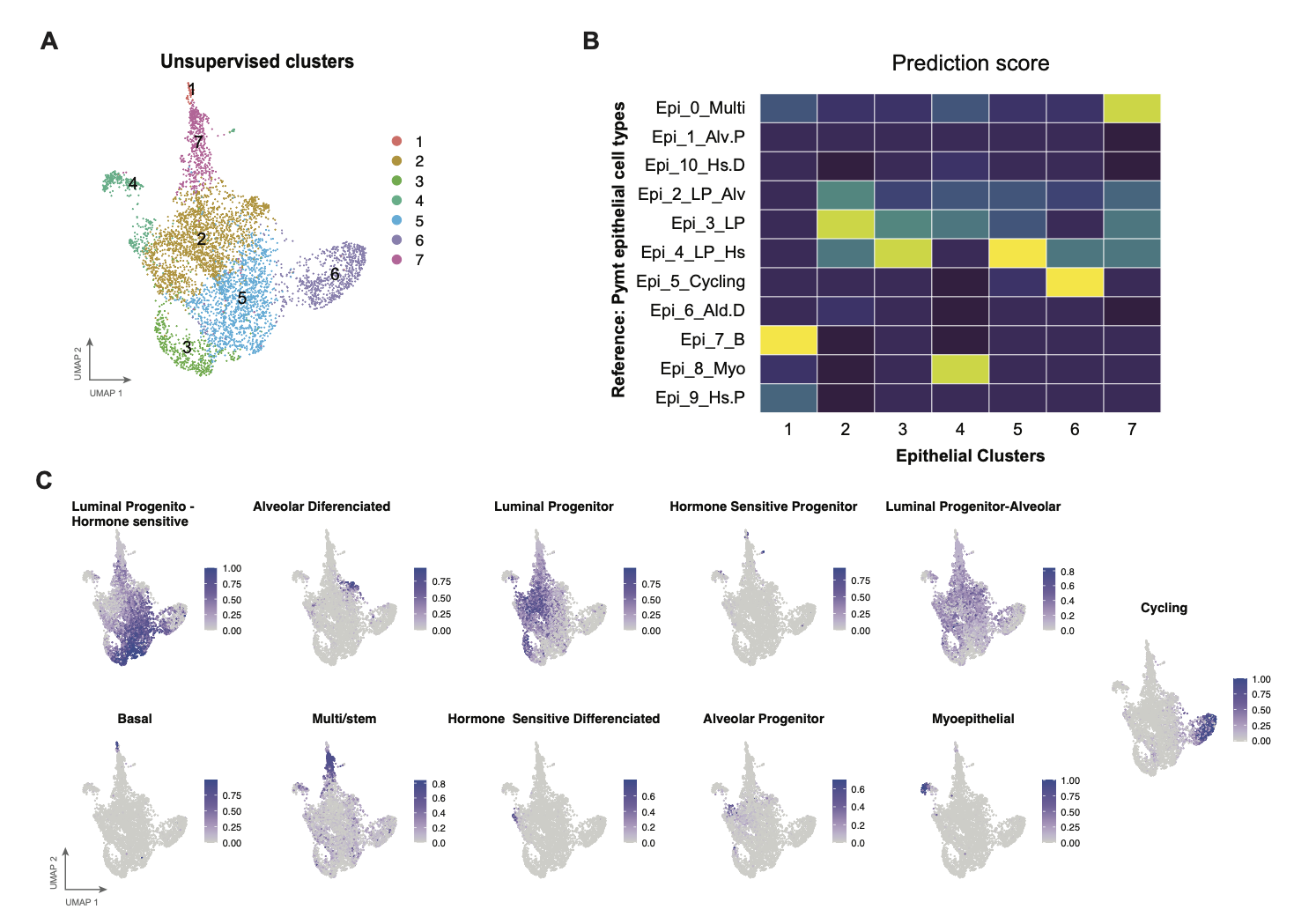

**Figure S3: A.** UMAP plot of the unsupervised clusters (resolution 0.7) of the epithelial cells. **B.** Predicted scores for epithelial subtypes from the PyMT reference on the unsupervised clusters (resolution 0.7) from the epithelial compartment. **C.** UMAP plots showing the prediction score per cell of each epithelial subtype from PyMT reference.

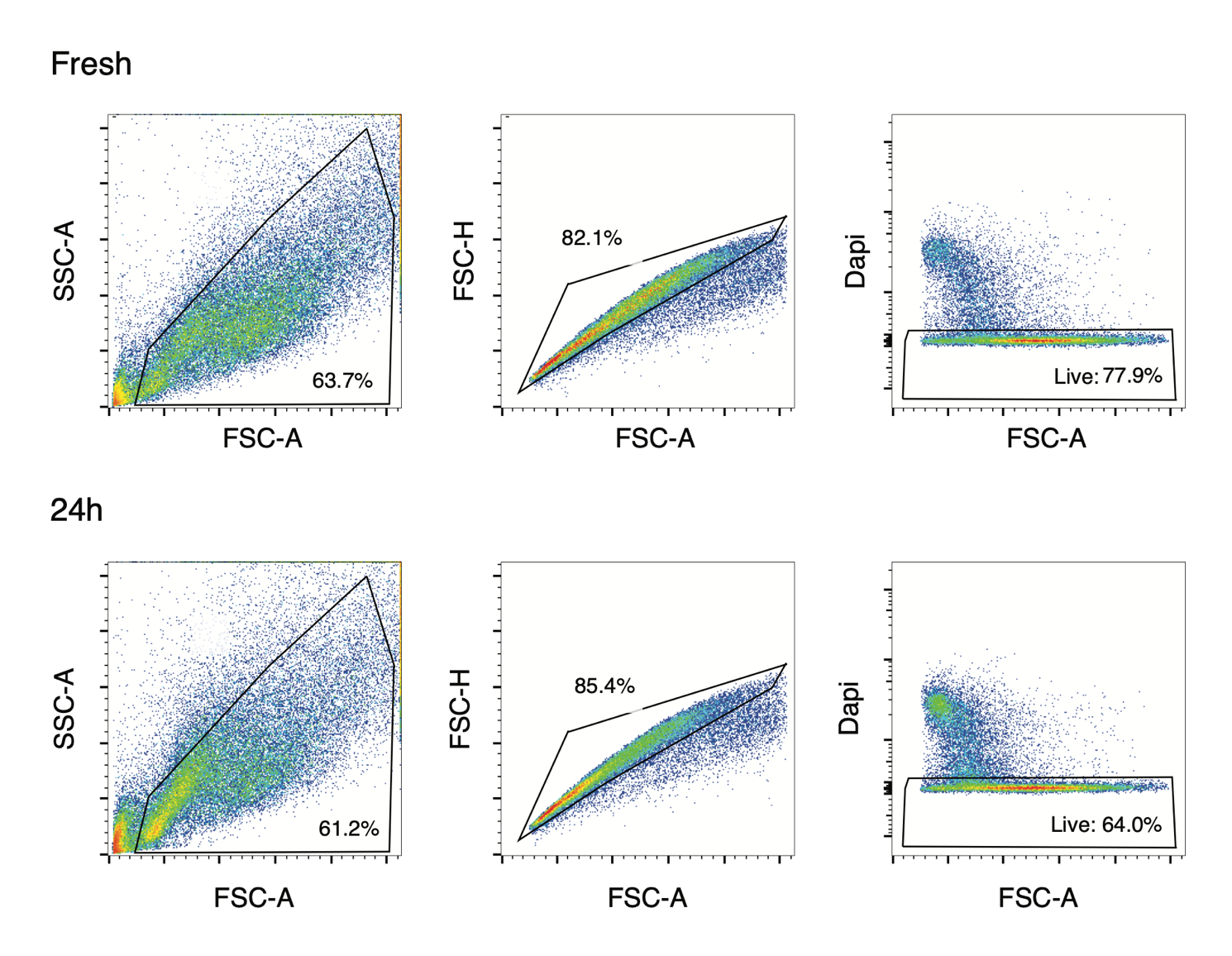

**Figure S4:** Flow cytometry gating strategy and representative comparison of the cell viability between cell suspensions from fresh tissue vs. after 24 hours at 4°C.

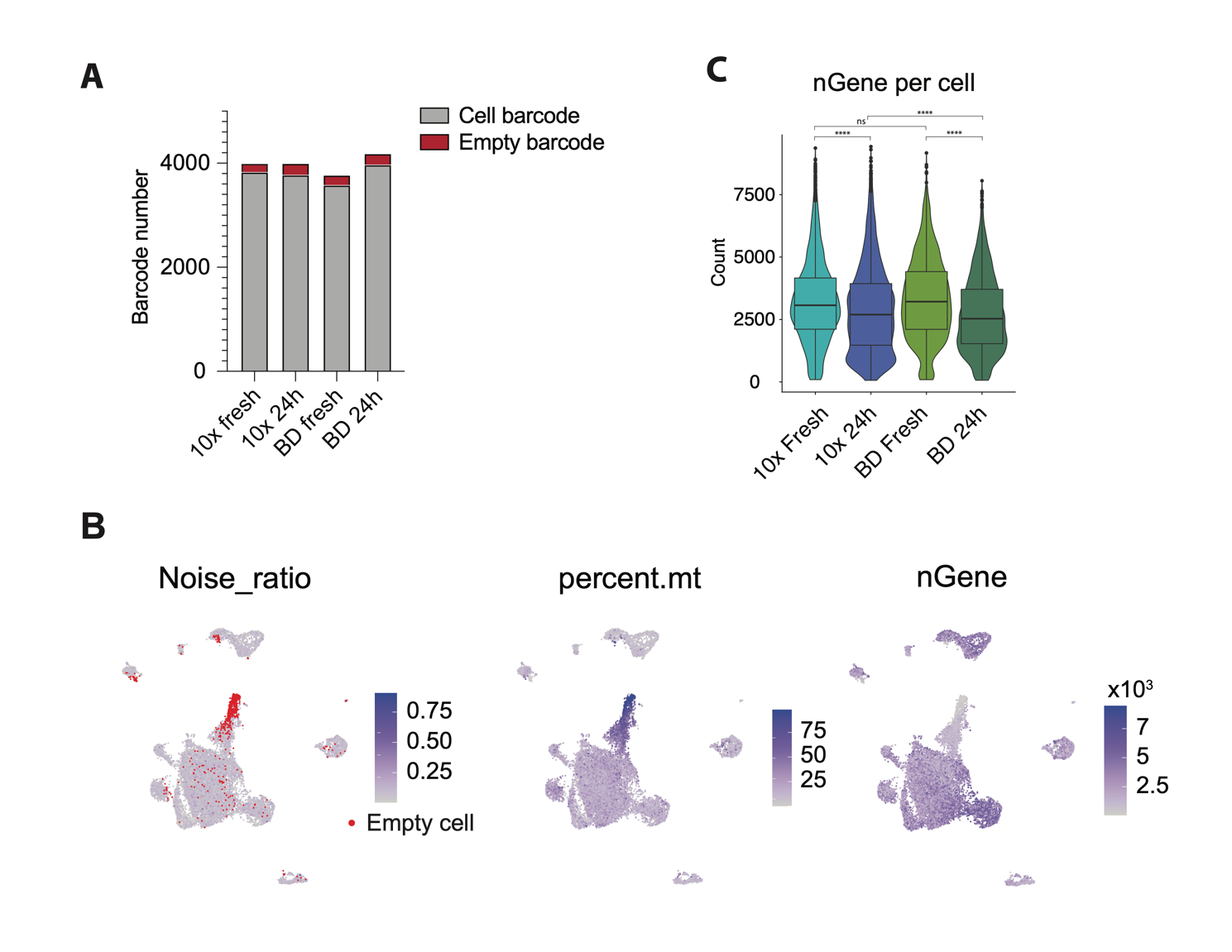

**Figure S5: A.** Number of barcodes assigned as putative cells or empty barcodes based on the transcriptome profile using the scAR package. **B.** UMAP visualizations of the noise ratio, percentages of mitochondrial genes (percent.mt) and the number of genes (nGene) detected. Cells marked in red are considered empty barcodes based on the scAR model. **C.** Comparison between the number of genes detected in 10x and BD Rhapsody of the fresh and damaged samples (24h).

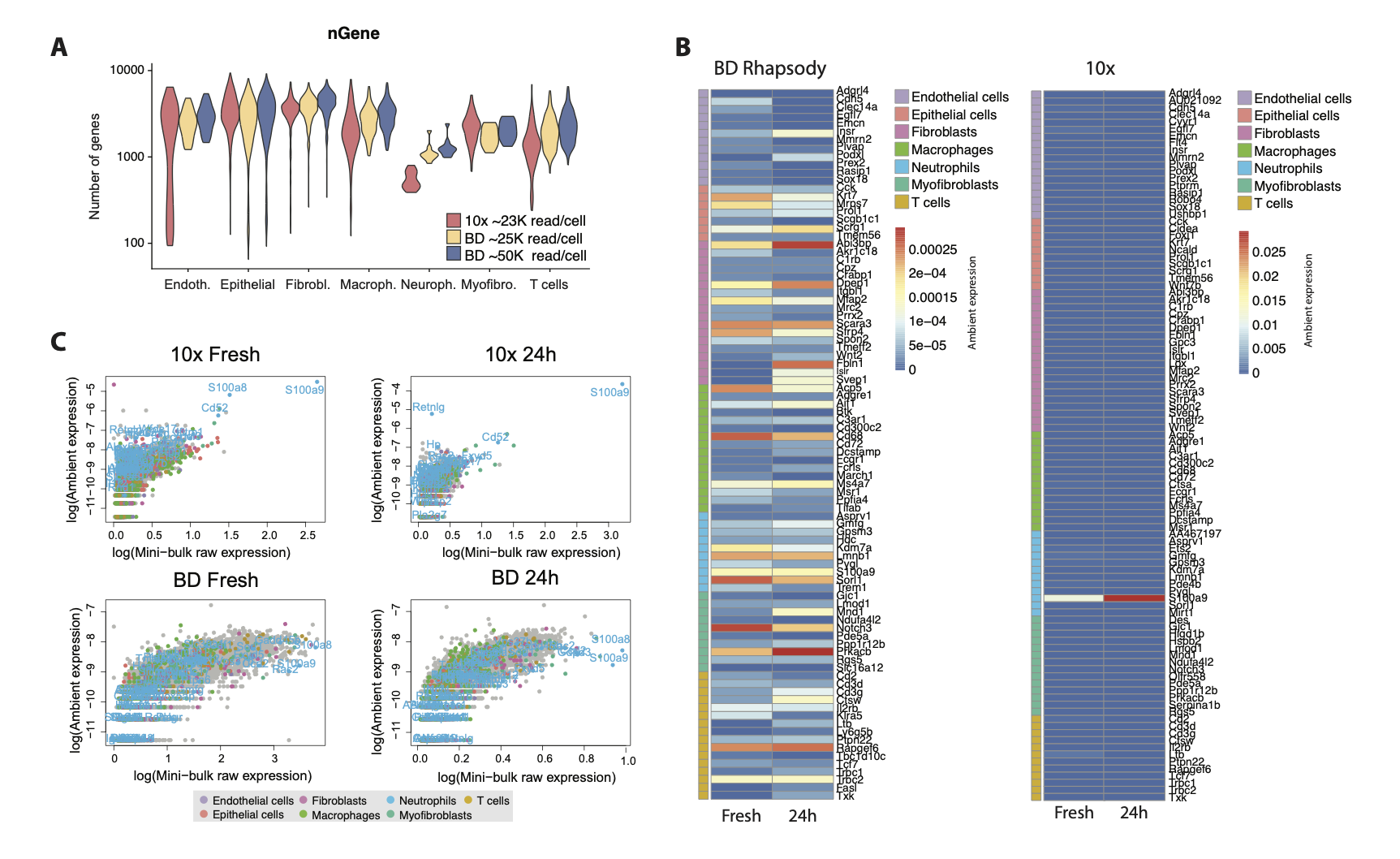

**Figure S6: A.** Violin plot showing the number of genes detected in each cell type in 10x Chromium and BD Rhapsody after downsampling the sequencing depth to obtain similar reads per cell than 10x.Y axis is shown logarithmic scale **B.** Heatmaps showing gene expression of the top markers of each cell type in the ambient pool. **C.** Scatter plot showing the correlation between the expression of each gene in the ambient pool (ambient expression) and the sum across single cells (mini-bulk raw expression). Genes are coloured based on the cell type they are markers for.
